# Variation in seed abundance predicts yolk fatty acid composition in a wild population of birds

**DOI:** 10.1101/2025.09.03.673935

**Authors:** Lucia Mentesana, Hong-Lei Wang, Michaela Hau, Caroline Isaksson, Martin N. Andersson

**Author notes:** **Author for correspondence:** Lucia Mentesana. These authors declare shared senior authorship (Caroline Isaksson, Martin N. Andersson).

## Abstract

Bird embryos develop inside eggs, which contain maternal substances that can shape offspring phenotype and fitness. Yolk fatty acids are a key energy source for the developing embryo, with omega-6 (ω-6) and omega-3 (ω-3) polyunsaturated fatty acids (PUFAs) obtained exclusively from the diet. Therefore, food availability during the breeding season is expected to influence yolk fatty acid composition and, consequently, embryo development and growth. However, the effects of dietary variation on yolk fatty acids remain unexplored in wild birds. We investigated inter-annual variation in yolk fatty acid composition of free-living great tits (*Parus major*) in relation to fluctuations in beech (*Fagus sylvatica*) fructification—their preferred food—across two years differing strongly in seed abundance. We hypothesized that differences in beech seed availability would alter yolk fatty acid composition between years of high and low fructification, with corresponding correlations to the fatty acid profiles of available seeds. Hence, we analyzed fatty acids from seeds of beech and two conifers (*Picea abies* and *Pinus sylvestris*), which account for most of the tree cover in the study area, and from 112 eggs collected from 107 nests over two years. Beech seeds contained higher proportions of saturated (SFA), monounsaturated (MUFA), and ω-3 PUFAs, and lower proportions of ω-6 PUFAs than conifer seeds. Correspondingly, egg yolks had more SFA, MUFA, and ω-3 PUFAs in the year of high beech abundance, and more ω-6 PUFAs in the year of low abundance. Notably, the proportion of the conifer-associated ω-6 PUFA pinolenic acid was 64.25 times higher in years of low compared to high beech fructification, suggesting marked inter-annual variation in yolk fatty acid composition, likely reflecting shifts in maternal diet. Future experiments need to establish causal links between diet and yolk composition and assess the fitness consequences of different seed resources for breeding females and their offspring.

## Introduction

Bird embryos develop inside an enclosed package of maternal substances in the egg yolk, including hormones, antioxidants, immune components, and nutrients (e.g., Saino et al. 2002, Tilgar et al. 2005, Mentesana et al. 2019a, Valcu et al. 2019). Variation in maternal deposition of these substances into the yolk has been shown to affect offspring phenotype and fitness (Saino et al. 2002, Parolini et al. 2017, Watson et al. 2018, Mentesana et al. 2021, 2025). Among these nutrients, fatty acids are an essential energy source for the growing embryo (Noble and Cocchi 1990), and certain omega-6 (ω-6) and ω-3 polyunsaturated fatty acids (PUFAs) are crucial for proper development and function of specific tissues and organs. One example is the ω-6 PUFA arachidonic acid (20:4n-6), which is important for brain function and the contractions of heart muscle cells (Hohl and Rösen 1987, Katsuki and Okuda 1995, Pavoine et al. 1999), in addition to its role in innate immune responses, where it is converted into pro-inflammatory eicosanoids. A high intake of ω-6 PUFAs relative to ω-3 PUFAs may exacerbate inflammatory responses, whereas a low ω-6/ω-3 ratio can have anti-inflammatory effects (Larsson et al. 2004, Cherian 2007, Gomes et al. 2012). Furthermore, the excitable membranes of the retina, brain, and skeletal muscle require a high proportion of the ω-3 PUFA docosahexaenoic acid (DHA; 22:6n-3) for proper functioning (Neuringer et al. 1988, Anderson et al. 1989, Speake et al. 1996, Mitchell et al. 1998, Salem Jr et al. 2001, Speake and Wood 2005).

Maternal deposition of substances into the egg yolk is influenced not only by female age and genetics (e.g., Ruuskanen et al. 2016, Valcu et al. 2019) but also by external factors such as ambient temperature and female nutrition (Stadelman and Pratt 1989, Noble and Cocchi 1990, Noble et al. 1996, Leskanich and Noble 1997, Surai et al. 2001, Twining et al. 2018). In particular, saturated (SFAs) and monounsaturated fatty acids (MUFAs) can be synthesized *de novo* by birds, whereas longer-chain ω-6 and ω-3 PUFAs can only be produced from strictly dietary precursors —linoleic- and α-linolenic acid, respectively (Larsson et al. 2004, Twining et al. 2016). Consequently, the composition of these fatty acids in the egg yolk is expected to be strongly influenced by the mother’s diet (Maldjian et al. 1996, Noble et al. 1996, Speake et al. 1999a, b, Surai et al. 2001, Surai and Speake 2008). Supporting this, robust data from poultry studies demonstrate a strong correlation between the fatty acid content of female ingested food and that found in egg yolk (e.g., Lin et al. 1991, Poureslami et al. 2012, Kralik et al. 2021).

In free-living bird species, diet can vary substantially across both temporal and spatial scales, affecting the fatty acid composition of their tissues (Conway et al. 1994, Egeler and Williams 2000, Pierce and McWilliams 2005, Andersson et al. 2015a, Isaksson et al. 2017). For example, the plasma fatty acid composition of great tits (*Parus major*) from northern Europe changes markedly between winter and the breeding season, reflecting a dietary shift from granivory in winter to insectivory in the warmer months (Andersson et al. 2015b). Furthermore, differences in fatty acid composition of both adult birds and egg yolks have been observed among populations of several passerine species, including great tits and the closely related blue tit, *Cyanistes caeruleus* (Bourgault et al. 2007, Andersson et al. 2015b, Toledo et al. 2016, Isaksson et al. 2017). Diet availability or dietary preferences during a given season may also vary between years, influencing the fatty acid intake during periods such as the breeding season. In granivorous birds, annual variation in tree fructification can have substantial effects on fatty acid intake, as different seeds have distinct fatty acid profiles (Beare-Rogers et al. 2001). This, in turn, may affect the fatty acids available for deposition in the egg yolk, potentially influencing yolk composition and, consequently, embryo development and growth. To our knowledge, the effects of yearly variation in diet availability on egg yolk fatty acid composition have not yet been investigated in a wild bird population.

Here, we investigated inter-annual variation in the fatty acid composition of great tit egg yolks in relation to fluctuations in the fructification of beech (*Fagus* sp.) trees. Beech seeds are a preferred food source for adult great tits during autumn and winter and strongly influence their survival (Perdeck et al. 2000). However, seed production varies greatly between years, with some years yielding abundant seeds and others producing few or none. The tit population under study inhabits an area where both beech trees and conifers (e.g., *Pinus* and *Picea* sp.) are abundant, and the birds have been observed consuming conifer seeds when beech seeds are scarce or unavailable (e.g., Sjöberg et al. 2007). We therefore hypothesized that differences in beech seed abundance would affect yolk fatty acid composition between years of high and low beech fructification, with corresponding correlations to the fatty acid profiles of the ingested seeds. Based on previously reported fatty acid compositions of different seeds (Prasad and Gülz 1989, Wolff and Bayard 1995, Vanhanen et al. 2017), we predicted that SFAs, MUFAs, and ω-3 PUFAs would be present in higher relative amounts in yolk during years when beech seeds were abundant. In contrast, conifer seeds contain higher levels of ω-6 PUFAs than beech seeds, including conifer-specific fatty acids such as pinolenic, taxoleic, and sciadonic acids (Xie et al. 2016, Vanhanen et al. 2017). Thus, in years with low beech seed availability, we predicted that a diet shift toward conifer seeds would result in higher relative yolk levels of ω-6 PUFAs, including the characteristic conifer seed fatty acids.

## Materials and methods

### Field site

We carried out this study in a nest-box population of great tits in the Dellinger Buchet, in Southern Germany (Bavaria; 48° 03′ N, 11° 13′ E, 620 m.a.s.l.). The Dellinger Buchet is a mixed forest with a mosaic of deciduous and coniferous trees, dominated by spruce (*Picea* sp.), beech (*Fagus* sp.), ash (*Fraxinus excelsior*), and European larch (*Larix decidua*), which make up 28.31%, 28.08%, 14.89%, and 12.97% of the forest, respectively (pers. comm. with the forester). This forest contains 125 nest boxes distributed over ∼1.30 km^2^, placed on tree trunks at 1.5-2.0 m height and spaced ∼40 m apart. We collected data on ambient temperature (°C) every hour from 12 i-buttons (DS9093A+ Thermochron iButton) placed at different locations within the forest.

### Beech tree fructification

We obtained information on the annual percentage of beech (*Fagus* sp.) mast fructification from the Bavarian State Minister for Food, Agriculture and Forestry (Bayerisches Staatsministerium für Ernährung, Landwirtschaft und Forsten 2018), and extracted the values using the R package ‘metaDigitise’ (Pick et al. 2019).

### Seed collection

We obtained tree seeds from the Bavarian Office for Forest Seeding and Planting, including from Norway spruce (*Picea abies*) and beech (*Fagus sylvatica*), which together account for 56.39% of the total tree coverage in our study population, as well as Scots pine (*Pinus sylvestris*). Scots pine seeds were included because great tits have been recorded feeding on its seeds (e.g., Gibb and Betts 1963), and although it is not present in the forest with our nest boxes, it is the second most abundant tree species in the surrounding area (Bayerisches Staatsministerium für Ernährung, Landwirtschaft und Forsten, 2018). Therefore, great tits might still utilize Scots pine seeds from nearby forests.

### Egg collection

Female great tits generally lay one egg per day. In our population, the mean clutch size is (mean ± SD) 8.45 ± 1.13 eggs. During the 2015 and 2016 breeding seasons, we checked each nest box every other day from the start of the breeding season. Once egg laying started, we marked each egg with a pencil to identify its position in the laying order. Great tit females vary substantially in the mean allocation of yolk components and in their plasticity along the laying sequence (e.g., Lessells et al. 2016, Mentesana et al. 2019). However, in our population, yolk fatty acids are highly repeatable within a female (*R* > 0.5 for SFAs and MUFAs and *R* > 0.9 for PUFAs; Mentesana et al. 2019). These repeatability estimates suggest that the middle egg of each clutch represents the average yolk content for a given clutch. We therefore collected the fourth egg from first and second clutches. We collected eggs between 8:00 and 13:00 h on the day they were laid and replaced them with a dummy egg. In total, we collected 112 eggs (N _*2015*_ = 45: 37 from first and 8 from second clutches; N _*2016*_ = 67: 53 from first and 14 from second clutches) from 107 nests (N _*2015*_ = 44, N _*2016*_ = 63) over the two years.

On the day of collection, we weighed and opened each freshly laid egg in the laboratory and separated the yolk from the albumen by rolling it on a piece of paper. We then homogenized the yolk by mixing it with an equal amount of distilled water (1 μl per mg of yolk) and stored it at −80 °C until further analysis.

### Seeds analysis

We analyzed fatty acid of seeds from three tree species (*F. sylvatica*., n = 4; *P. abies*, n = 3; *P. sylvestris*, n = 3). For *P. abies* and *P. sylvestris*, which have comparatively small seeds, we used whole seeds (approximately 8-10 mg and 6-7 mg, respectively) for fatty acid extractions. For the larger beech seeds, we removed the seed coat, cut the seeds into smaller pieces, and combined several random pieces from each seed (totaling 17–25 mg) for each extraction. We carried out acid methanolysis using 1 mL of 2% H_2_SO_4_ in methanol. We kept the samples at 90[°C, and after 20 min, we crushed the seed material using a glass rod. The reaction then proceeded for another 40 min, after which we added 1 mL deionized water and 1 mL heptane. We thoroughly vortexed the samples, transferred the heptane layer containing the extracted FAMEs to a new vial, and washed it twice with 1 mL of H_2_O. We removed residual water using anhydrous sodium sulfate.

We analyzed all fatty acids extracts using an Agilent 5975 mass spectrometer (MS) coupled to an Agilent 6890 gas chromatograph (GC) with an HP-INNOWax PEG column (30 m × 0.25 mm i.d., 0.25 mm film thickness; Agilent). We performed analyses and quantification of chromatograms using ChemStation software (Agilent). Finally, we identified FAMEs by comparing mass spectra and retention times with those of synthetic standards (Supelco 37-Component FAME Mix, Sigma-Aldrich).

### Egg yolk analysis

We also extracted fatty acids from 112 egg yolks following previously described methods (Toledo et al. 2016, Mentesana et al. 2019, 2021). We extracted total lipids from approximately 5 mg of yolk using 50 μl chloroform and methanol (2:1 v/v) mixed with 16.65 μg of the internal standard methyl *cis*-10-heptadecenoate (>99%; Aldrich). Then, we dried the samples under N_2_ and carried out base methanolysis (using 100 mL 0.5 M KOH/Me; reaction proceeded for 1h at 40 °C and terminated using 100 mL 0.5 M HCl/Me) to transform the fatty acids into the corresponding fatty acid methyl esters (FAMEs). Next, we extracted the FAMEs with heptane (> 99%; VWR Prolabo), washed the extracts with deionized H_2_O, and removed residual water using anhydrous sodium sulfate. The GC-MS analysis and FAME identification were performed as described above for the seeds.

### Data handling and statistical analysis

To statistically analyze the fatty acids in seeds and egg yolks, we first calculated the proportion of each fatty acid by dividing its peak area of each by the sum of the peak areas of all fatty acids in each individual sample (following Andersson et al. 2015, Isaksson et al. 2017). We then combined the proportions of all individual fatty acids within a certain chemical class of fatty acid to obtain relative levels of total SFA, MUFA, ω-6 and ω-3 PUFAs (Andersson et al. 2015b, Isaksson et al. 2017).

Second, to study differences in seed fatty acid proportions, we ran five univariate linear models, each with the logit-transformed proportion (Warton and Hui 2011) of a specific fatty acid or the omega-6 to omega-3 ratio as the response variable, and ‘tree species’ as a fixed factor (levels: beech, spruce, pine).

Third, to study the differences in yolk fatty acid proportions between years, we ran five univariate linear mixed-effects models. We fitted the logit-transformed proportion of each fatty acid and the total omega-6 to total omega-3 PUFA ratio as response variables in separate models, each including ‘year’ (levels: 2015, 2016) as a fixed factor. We also fitted ‘date’ when each egg was collected and ‘mean ambient temperature’ for the three days prior to the lay date of each egg as covariates. Both covariates were mean-centered (i.e., mean value = 0, standard) because they differed in their scales of magnitude. We also included ‘nest ID’ and ‘female ID’ as random factors in the model testing differences in SFA between years. For the remaining four models, we included only ‘nest ID’ because ‘female ID’ explained no variance.

We performed all our analyses using the R-packages “*lme4*” and “*arm*” in R-3.3.3 (R Core Team 2013) in a Bayesian framework with non-informative priors. We assumed a Gaussian error distribution, which was confirmed for all response variables after visual inspection of model residuals. We based model structure on the study question and the biology of the species rather than on model selection (Korner-Nievergelt et al. 2015). We used the “*sim*” function to simulate posterior distributions of the model parameters. From 10,000 simulations, we extracted the 95% Bayesian credible interval (CrI) around the mean (Gelman and Hill 2007) and assessed statistical support by obtaining the posterior distribution of each parameter. We considered an effect to be ‘statistically meaningful’ when the estimated CrI did not include zero (Korner-Nievergelt et al. 2015).

## Results

### Beech tree fructification

Beech tree fructification was considerably higher in 2015 (23.31%) than in 2016 (1.01%; mean fructification over 21 years ± SD = 11.26 ± 2.79; Bayerisches Staatsministerium für Ernährung, Landwirtschaft und Forsten 2018).

### Fatty acids in tree seeds

The proportion of all four fatty acid groups —SFA, MUFA, ω-6 and ω-3 PUFAs —differed among the three tree species (Figure 1A-D; Supplementary Table 1). Beech (*F. sylvatica*) seeds, the preferred food source of great tits, contained higher proportions of SFA, MUFA, and ω-3 PUFAs, and a lower proportion of ω-6 PUFAs compared to the spruce (*P. abies*) and pine (*P. sylvestris*) seeds. The overall fatty acid composition was similar in seeds from the two conifer species. The ratio of total ω-6 to total ω-3 PUFAs was lower in beech seeds compared to spruce and pine seeds (Figure 1E; Supplementary Table 1).

**Figure 1:**
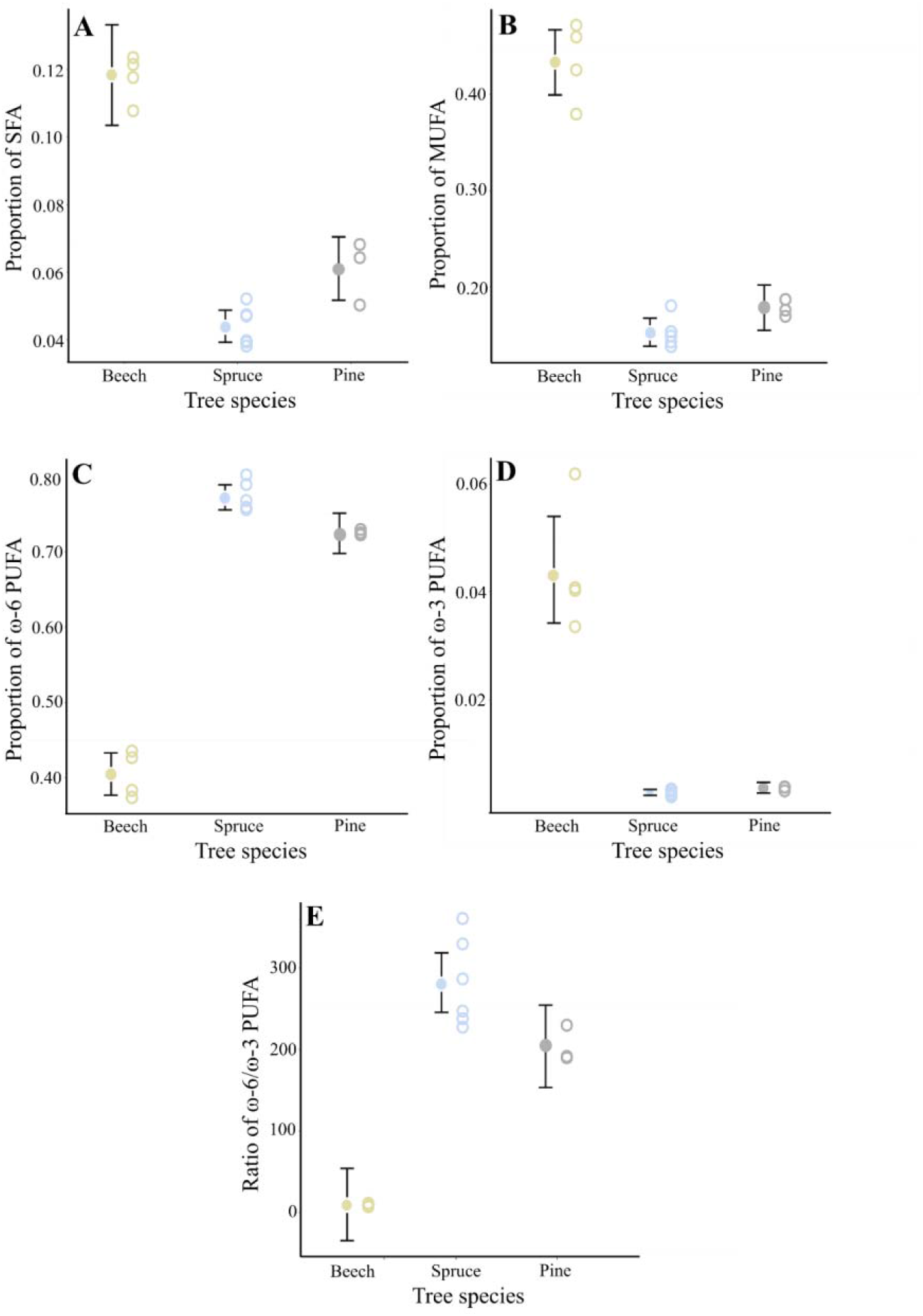
Proportion of total A) saturated (SFA), B) monounsaturated (MUFA), C) ω-6 polyunsaturated (PUFA), (D) ω-3 PUFA fatty acids, and (E) the ratio between the two PUFA classes in seeds of beech (*Fagus sylvatica*), spruce (*Picea abis*), and pine (*Pinus sylvestris*) trees. Yellow, light blue and grey open circles show the raw data of each tree species, respectively. Model mean estimates as shown as yellow, light blue and grey filled circles, with 95% credible intervals indicated by black vertical bars. A statistically significant difference between groups can be inferred if the credible interval of one group does not overlap with the mean estimate of the other group.

Differences in the mean percentages of the four fatty acid groups were driven both by variations in individual fatty acids and by the presence of certain fatty acids unique to certain tree species. Specifically, one SFA (behenic acid) and one MUFA (erucic acid) were detected only in beech seeds, whereas four PUFAs (pinolenic-, sciadonic-, taxoleic-, and eicosadienoic acids) were detected exclusively in the conifer seeds (Table 1). Taxoleic acid and one of the detected eicosadienoic acids are ω-9 PUFAs, biosynthesized along the same pathway as the ω-6 PUFAs pinolenic- and sciadonic acid.

**Table 1:**
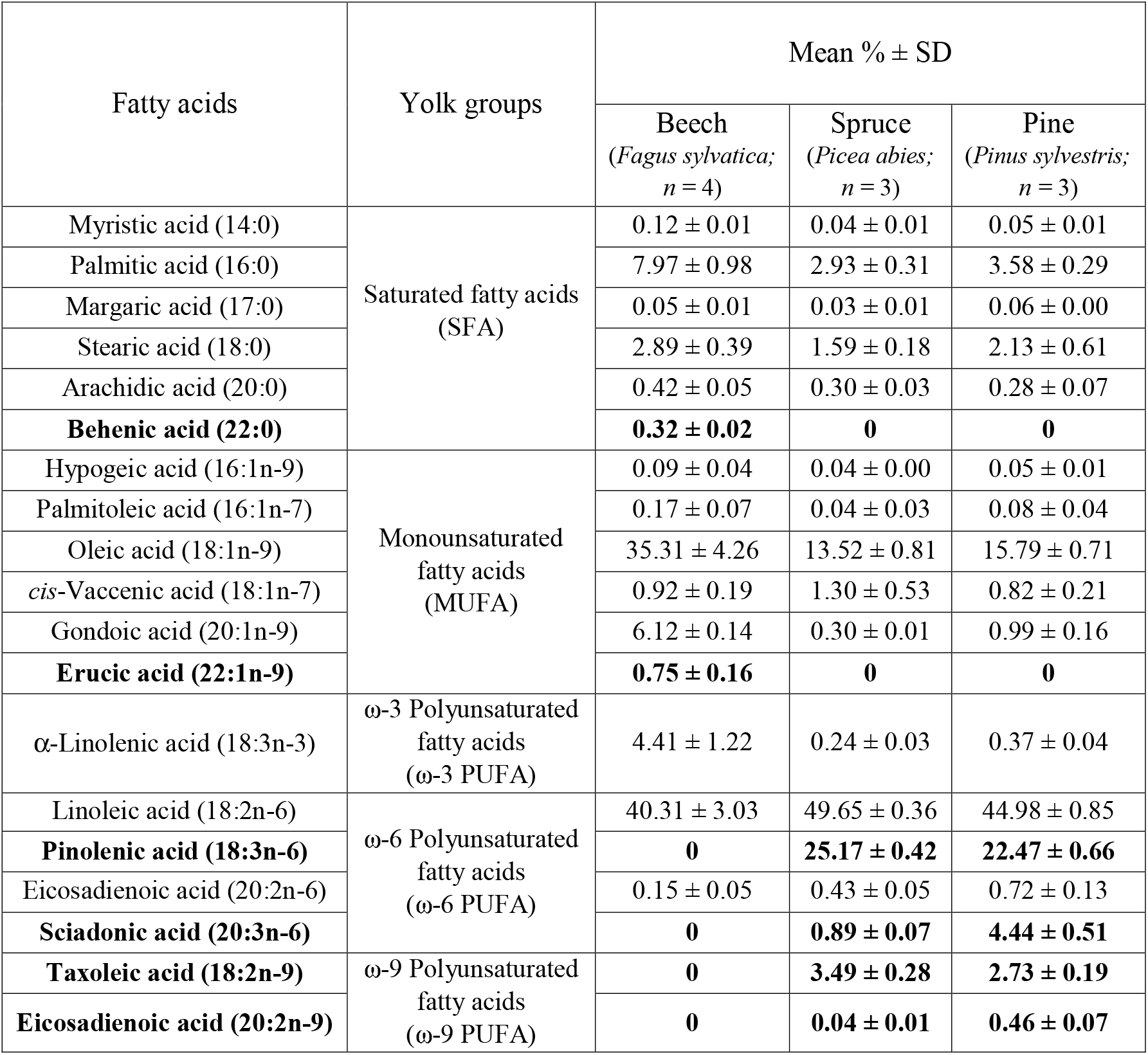
Overall relative abundance (% of total fatty acid content) of fatty acids present in the seeds of three tree species (beech: *Fagus sylvatica*, spruce: *Picea abies* and pine: *Pinus sylvestris*). For fatty acids lacking a trivial name, we provide their systematic names. Number of carbon atoms:number of double bonds and position are written between brackets. Highlighted in bold are those fatty acids present in the seeds of only beech or only the conifer (i.e., spruce and pine) species.

### Fatty acids in egg yolks

In great tit egg yolks, the proportion of all four fatty acid groups – SFA, MUFA, ω-6 and ω-3 PUFAs – differed between the two years of study (Figure 2A-D; Supplementary Table 2). The yolks collected in the year with high beech seed abundance (2015) had higher proportions of SFA, MUFA, and ω-3 PUFAs, and a lower proportion of ω-6 PUFAs compared to yolks collected in the year with low beech seed availability (2016). Accordingly, the ratio between ω-6 and ω-3 yolk PUFAs was lower in 2015 than in 2016 (Figure 2E; Supplementary Table 2).

**Figure 2:**
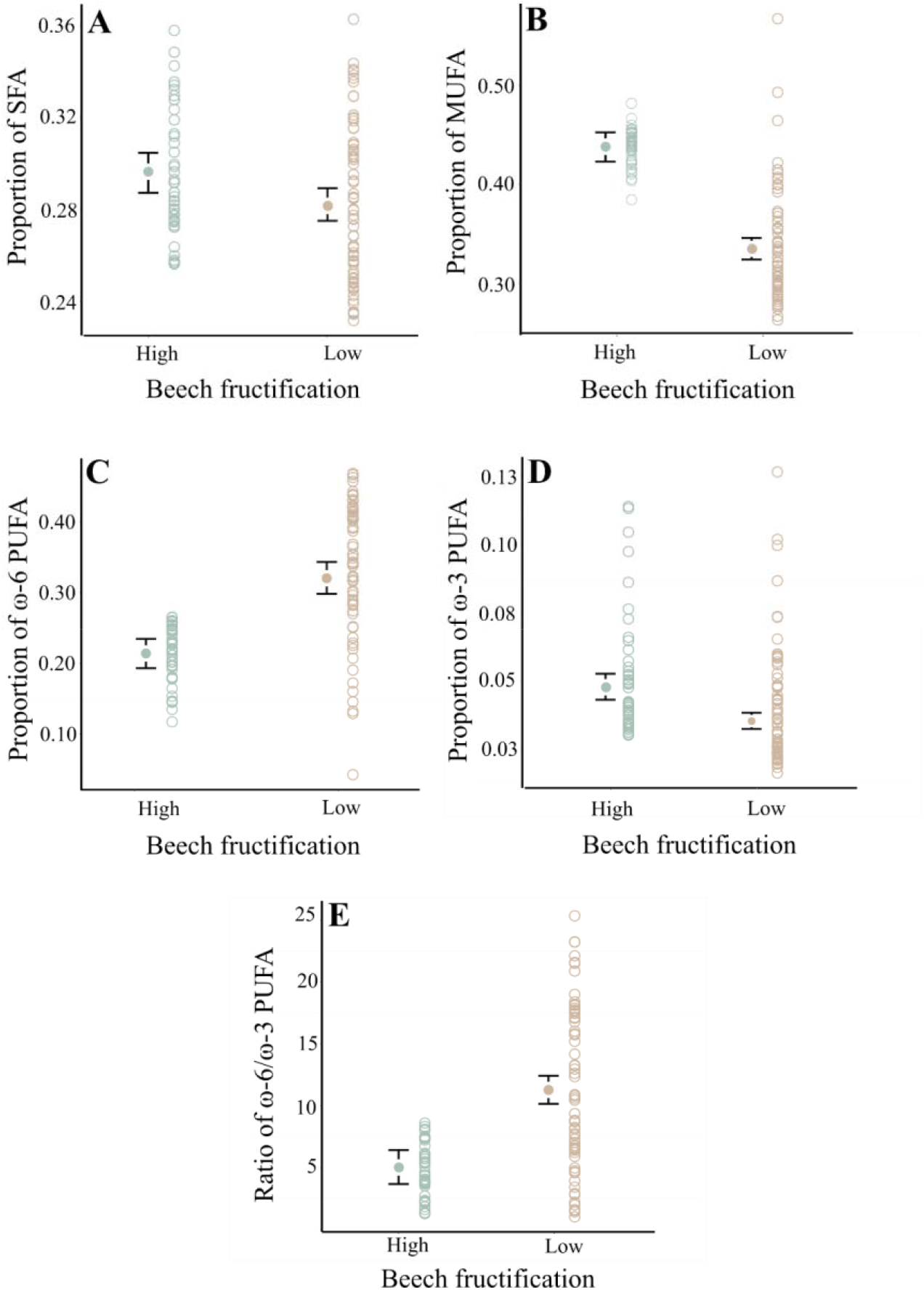
Proportion of total A) saturated (SFA), B) monounsaturated (MUFA), C) ω-6 polyunsaturated (PUFA), (D) ω-3 PUFA fatty acids, and (E) the ratio between the two PUFA classes in great tit egg yolk during two years with either high (2015) or low (2016) availability of beech seeds. Green and orange open circles show the raw data for eggs collected in 2015 and 2016, respectively. Model mean estimates as shown as green and orange filled circles, with 95% credible intervals indicated by black vertical bars. A statistically significant difference between groups can be inferred if the credible interval of one group does not overlap with the mean estimate of the other group.

The difference in fatty acid group proportions observed between years was generally driven by small differences in the proportion of individual fatty acids. However, certain fatty acids showed dramatic year-to-year differences; in particular, the proportions of the conifer seed-associated ω-6 PUFA pinolenic acid was 64.25x higher in 2016, when beech fructification was low, compared to 2015 (Table 2; Supplementary Figure 1). The proportion two other conifer seed-associated PUFAs, sciadonic and taxoleic acid, also displayed large increases (9.0x and 8.67x, respectively) from 2015 to 2016. Yet, it should be noted that the relative proportion of these latter PUFAs are still small.

**Table 2:** Overall relative abundance (% of total fatty acid content) of fatty acids present in great tit egg yolks. For fatty acids without a trivial name, we provide the systematic names. Number of carbon atoms:number of double bonds and position are written within brackets. Note that the proportion of most fatty acids changed between years, with some showing large differences (e.g., pinoleic acid).

| Yolk fatty acid | Fatty acid class | Relative abundance<br>(mean % ± SD; $n_{total} = 112$) | |
| --- | --- | --- | --- |
| | | 2015 ($n = 45$) | 2016 ($n = 67$) |
| Lauric acid (12:0) | Saturated fatty acids (SFA) | 0.02 ± 0.02 | 0.02 ± 0.02 |
| Myristic acid (14:0) | | 0.55 ± 0.36 | 0.44 ± 0.25 |
| Pentadecanoic acid (15:0) | | 0.09 ± 0.02 | 0.10 ± 0.05 |
| Palmitic acid (16:0) | | 20.15 ± 2.67 | 18.95 ± 3.45 |
| Margaric acid (17:0) | | 0.57 ± 0.13 | 0.57 ± 0.13 |
| Stearic acid (18:0) | | 8.09 ± 0.61 | 8.28 ± 0.72 |
| Hypogeic acid (16:1n-9) | Monounsaturated fatty acids (MUFA) | 1.35 ± 0.24 | 1.08 ± 0.24 |
| Palmitoleic acid (16:1n-7) | | 1.83 ± 1.38 | 1.29 ± 0.61 |
| Oleic acid (18:1n-9) | | 37.61 ± 2.32 | 28.75 ± 5.35 |
| *cis*-Vaccenic acid (18:1n-7) | | 1.62 ± 0.51 | 1.96 ± 0.33 |
| Gondoic acid (20:1n-9) | | 0.06 ± 0.02 | 0.06 ± 0.02 |
| Paullinic acid (20:1n-7) | | 1.36 ± 0.66 | 0.36 ± 0.09 |
| Hexadecadienoic acid (16:2n-6) | ω-6 Polyunsaturated fatty acids (ω-6 PUFA) | 0.05 ± 0.02 | 0.14 ± 0.07 |
| γ-Linolenic acid (18:3n-6) | | 0.11 ± 0.02 | 0.15 ± 0.04 |
| Linoleic acid (18:2n-6) | | 18.46 ± 3.59 | 26.77 ± 7.42 |
| Pinolenic acid (18:3n-6) | | 0.04 ± 0.11 | 2.57 ± 1.42 |
| Eicosadienoic acid (20:2n-6) | | 0.32 ± 0.08 | 0.58 ± 0.21 |
| dihomo-γ-Linolenic acid (20:3n-6) | | 0.32 ± 0.06 | 0.72 ± 0.29 |
| Sciadonic acid (20:3n-6) | | 0.03 ± 0.06 | 0.27 ± 0.12 |
| Arachidonic acid (20:4n-6) | | 1.96 ± 0.28 | 2.19 ± 0.53 |
| Adrenic acid (22:4n-6) | | 0.23 ± 0.08 | 0.31 ± 0.09 |
| α-Linolenic acid (18:3n-3) | ω-3 Polyunsaturated fatty acids (ω-3 PUFA) | 2.89 ± 1.79 | 2.01 ± 1.69 |
| Eicosapentaenoic acid (20:5n-3) | | 0.33 ± 0.10 | 0.32 ± 0.23 |
| Docosapentaenoic acid (22:5n-3) | | 1.63 ± 0.37 | 1.37 ± 0.49 |
| Docosahexaenoic acid (22:6n-3) | | 0.28 ± 0.06 | 0.22 ± 0.13 |
| Taxoleic acid (18:2n-9) | ω-9 Polyunsaturated fatty acids (ω-9 PUFA) | 0.06 ± 0.02 | 0.52 ± 0.28 |

Finally, the proportion of SFA, MUFA, ω-3 PUFAs increased, while the proportion of ω-6 PUFAs decreased over the breeding season (Supplementary Table 2). In addition, ω-3 PUFA proportions increased with rising temperature (Supplementary Table 2).

## Discussion

Our results show significant inter-annual variation in the relative abundances of yolk SFA, MUFA, and ω-3 and ω-6 PUFAs, as well as in the ω-6/ω-3 PUFA ratio. This variation likely reflects yearly differences in maternal diet: in years of local beech seed abundance, females probably consume more of this preferred food item, whereas in years of beech scarcity, a greater intake of conifer seeds is suggested by the presence of ‘conifer-diagnostic’ fatty acids in the yolk (primarily pinolenic acid, but also sciadonic- and taxoleic acid; Xie et al. 2016). These findings suggest that the maternal diet of wild birds shapes the fatty acid composition of egg yolks, which may, in turn, influence offspring growth, phenotype, and fitness across years (Mentesana et al. 2021).

Compared to the long-term mean beech seed fructification (11%), 2015 showed high fructification (23%), followed by very low fructification in 2016 (1%). This sharp reduction likely forced female great tits to switch from beech to conifer seeds. Seed analyses confirmed that beech contains higher proportions of SFA, MUFA, and ω-3 PUFA, and less ω-6 PUFA than conifers, consistent with previous studies (Prasad & Gülz, 1989; Wolff & Bayard, 1995; Vanhanen et al., 2017). Correspondingly, yolks contained more SFA, MUFA, and ω-3 PUFA in 2015, but more ω-6 PUFA in 2016, strongly altering the ω-6/ω-3 ratio. Since α-linolenic and linoleic acid are strictly dietary (Williams and Buck 2010) and maternal fatty acid plasma composition closely tracks yolk deposition (e.g., Lin et al. 1991, Poureslami et al. 2012, Kralik et al. 2021), the observed yolk profiles likely reflect this dietary shift. In line with the idea that changes in female diet influence yolk composition, independent of the year females laying eggs later in the breeding season had higher yolk ω-3 PUFA relative to ω-6 PUFA proportions (Supplementary Table 2). This pattern may reflect increasing caterpillar availability during the breeding season and a decreasing reliance on seeds, with caterpillars being rich in the ω-3 PUFA α-linolenic acid (Isaksson and Andersson 2007, Isaksson et al. 2015, Andersson et al. 2015a).

Great tits may prefer beech seeds because of their high nutritional value. In particular, the difference in ω-3 PUFA content between beech and conifer seeds is striking: our analyses showed that beech seeds contained about 4.5% ω-3 PUFAs, compared to only 0.3% in conifer seeds. Since ω-3 PUFAs are scarce in terrestrial food chains (Twining et al. 2016, Indykiewicz et al. 2023), the 4.5% found in beech seeds represents a substantial source of these essential nutrients, which are critical for the proper functioning of the retina, brain, and skeletal muscle membranes (e.g., Neuringer et al. 1988, Anderson et al. 1989, Maldjian et al. 1996, Speake et al. 1996, Mitchell et al. 1998, Salem Jr et al. 2001, Speake and Wood 2005). In contrast, conifer seeds contained an extremely high proportion of ω-6 PUFAs (70–80%), compared to around 40% in beech seeds. This is mainly due to the conifer-specific fatty acid pinolenic acid (Wolff and Bayard 1995) and, to a lesser extent, linoleic acid.

An increased allocation of ω-3 PUFAs into the yolk during beech seed rich years might have led to a higher reproductive success. A yolk composition of relatively more ω-3 to ω-6 has many advantages for the developing chicks, including enhanced neurodevelopment, visual acuity, immune balance, physiological resilience, muscle coordination and growth (Sujatha and Narahari 2011, Thanabalan and Kiarie 2021). Indeed, our previous study showed that great tit nests with eggs containing higher proportions of ω-3 PUFAs, as well as SFA and MUFAs (which also are abundant in beech seeds), were positively associated with higher numbers of hatchlings and fledglings than nests with eggs containing lower proportions of these fatty acids (Mentesana et al. 2021). In contrast, a diet composition that raises the yolk ω-6/ω-3 PUFA ratio, such as high intake of conifer seeds, may be less beneficial for the chicks since such a diet is expected to stimulate proinflammatory responses increasing the severity of an infection, and it also enhances the production of reactive oxygen species (ROS), which may cause oxidative damage (Larsson et al. 2004, Romieu et al. 2007, Isaksson 2015, Mentesana et al. 2021). In our study system, however, the increased proportion of ω-6 PUFA in yolk during the year with low beech fructification was partly driven by pinolenic acid, which has been suggested to promote health in broiler chickens via anti-inflammatory responses (e.g., Chen et al. 2015, Xie et al. 2016, Vienola et al. 2018). However, it remains unknown whether the high proportion of linolenic acid (the precursor of the pro-inflammatory arachidonic acid) in conifer seeds and the yolks from 2016 (Tables 1 & 2) outweighs the potential anti-inflammatory effects of pinolenic acid (Park et al., 2013; Xie et al., 2016). Future research is needed to determine the fitness consequences of beech and conifer seed components on breeding wild birds, as well as the specific effects of compounds such as pinolenic acid on avian physiology, health, and offspring development.

## Conclusions

In summary, our results show that yolk fatty acid composition varies markedly between years, reflecting differences in food availability. Because beech seeds are nutritionally rich (i.e., they contain high proportion of ω-3 PUFAs which are scarce for terrestrial birds), female great tits likely consume them preferentially whenever available, potentially enhancing their fitness. However, experimental studies are needed to test these hypotheses. For example, one could manipulate beech seed availability with artificial feeders across years in nearby forests. Female yolk deposition in these experimental forests could then be compared with that in control forests, where birds are exposed to natural seed fluctuations. Linking both control and experimental sites to reproductive success would provide a direct test of how diet availability influences yolk fatty acid composition and the potential fitness benefits of beech seed consumption.

---

**Supplementary Table 1:** Results from linear models explaining variation in the proportion of fatty acid content present in the seeds of three tree common species in the population. ‘Tree species’ (Beech: *Fagus sylvatica*, Spruce: *Picea abis* and Pine: *Pinus sylvestris*) was fitted as a fixed factor. We present fixed (β) parameters with their 95% credible intervals (CrI). Differences across tree species are statistically significant if zero is not included within the 95% CrI.

| | SFA<sup>a</sup> | MUFA<sup>a</sup> | ω-6 PUFA<sup>a</sup> | ω-3 PUFA<sup>a</sup> | ω-6 PUFA/<br>ω-3 PUFA |
| --- | --- | --- | --- | --- | --- |
| Fixed factors $\beta$ (95% CrI) | | | | | |
| Intercept | -2.016<br>(-2.157;<br>-1.875) | -0.269<br>(-0.408;<br>-0.132) | -0.388<br>(-0.507;<br>-0.271) | -3.102<br>(-3.345;<br>-2.855) | 9.118<br>(-34.961;<br>51.902) |
| Spruce<br>(*Picea abis*) | -1.071<br>(-1.253;<br>-0.886) | -1.446<br>(-1.621;<br>-1.269) | 1.615<br>(1.459;<br>1.769) | -2.776<br>(-3.092;<br>-2.466) | 272.151<br>(215.197;<br>328.583) |
| Pine<br>(*Pinus sylvestris*) | -0.726<br>(-0.939;<br>-0.509) | -1.267<br>(-1.477;<br>-1.052) | 1.364<br>(1.185;<br>1.546) | -2.527<br>(-2.904;<br>-2.157) | 194.271<br>(127.071;<br>261.447) |

<sup>a</sup> SFA, saturated fatty acids; MUFA, monounsaturated fatty acids; PUFA, polyunsaturated fatty acids. All fatty acid proportions were logit transformed.

**Supplementary Table 2:** Results from linear mixed-effects models explaining variation in yolk fatty acid proportion between years. ‘Year’ (2015 vs. 2016) was fitted as a fixed factor, ‘date’ when each egg was collected and ‘mean ambient temperature’ were fitted as covariates, and ‘female’ and ‘nest ID’ were fitted as random factors. All covariates were mean centered. We present fixed (β) and random (σ2) parameters with their 95% credible intervals (CrI). Fixed factors and covariates are statistically meaningful if zero is not included within the 95% CrI.

| | SFA<sup>a</sup> | MUFA<sup>a</sup> | ω-6 PUFA<sup>a</sup> | ω-3 PUFA<sup>a</sup> | ω-6 PUFA/<br>ω-3 PUFA |
| --- | --- | --- | --- | --- | --- |
| Fixed factors $\beta$ (95% CrI) | | | | | |
| Intercept | -0.866<br>(-0.908;<br>-0.825) | -0.249<br>(-0.308;<br>-0.189) | -1.305<br>(-1.428;<br>-1.183) | -2.995<br>(-3.099;<br>-2.889) | 5.067<br>(3.732;<br>6.423) |
| Year | -0.067<br>(-0.119;<br>-0.013) | -0.435<br>(-0.513;<br>-0.357) | 0.553<br>(0.394;<br>0.713) | -0.309<br>(-0.445;<br>-0.172) | 6.209<br>(4.457;<br>7.969) |
| Date<sup>b</sup> | 0.091<br>(0.059;<br>0.122) | 0.062<br>(0.017;<br>0.109) | -0.176<br>(-0.261;<br>-0.092) | 0.182<br>(0.092;<br>0.271) | -2.519<br>(-3.641;<br>-1.377) |
| Mean temperature<sup>c</sup> | 0.007<br>(-0.027;<br>0.042) | -0.022<br>(-0.072;<br>0.026) | -0.031<br>(-0.128;<br>0.064) | 0.123<br>(0.034;<br>0.209) | -0.572<br>(-1.716;<br>0.567) |
| Random factors $\sigma^2$ (95% CrI) | | | | | |
| Female ID | 0.003<br>(0.002; 0.004) | - | - | - | - |
| Nest ID | 0.012<br>(0.009; 0.016) | 0.024<br>(0.019;<br>0.030) | 0.144<br>(0.124;<br>0.169) | 0.042<br>(0.030;<br>0.057) | 8.471<br>(6.272;<br>11.267) |
| Residual variance | 0.002<br>(0.001; 0.002) | 0.016<br>(0.012;<br>0.021) | 0.025<br>(0.020;<br>0.033) | 0.085<br>(0.066;<br>0.113) | 12.233<br>(9.493;<br>16.289) |

<sup>a</sup> SFA, saturated fatty acids; MUFA, monounsaturated fatty acids; PUFA, polyunsaturated fatty acids.
All fatty acid proportions were logit transformed.
<sup>b</sup> Date when the fourth egg was collected.
<sup>c</sup> Mean ambient temperature for the 3 days prior to the lay date of each egg.

**Supplementary Figure 1:**
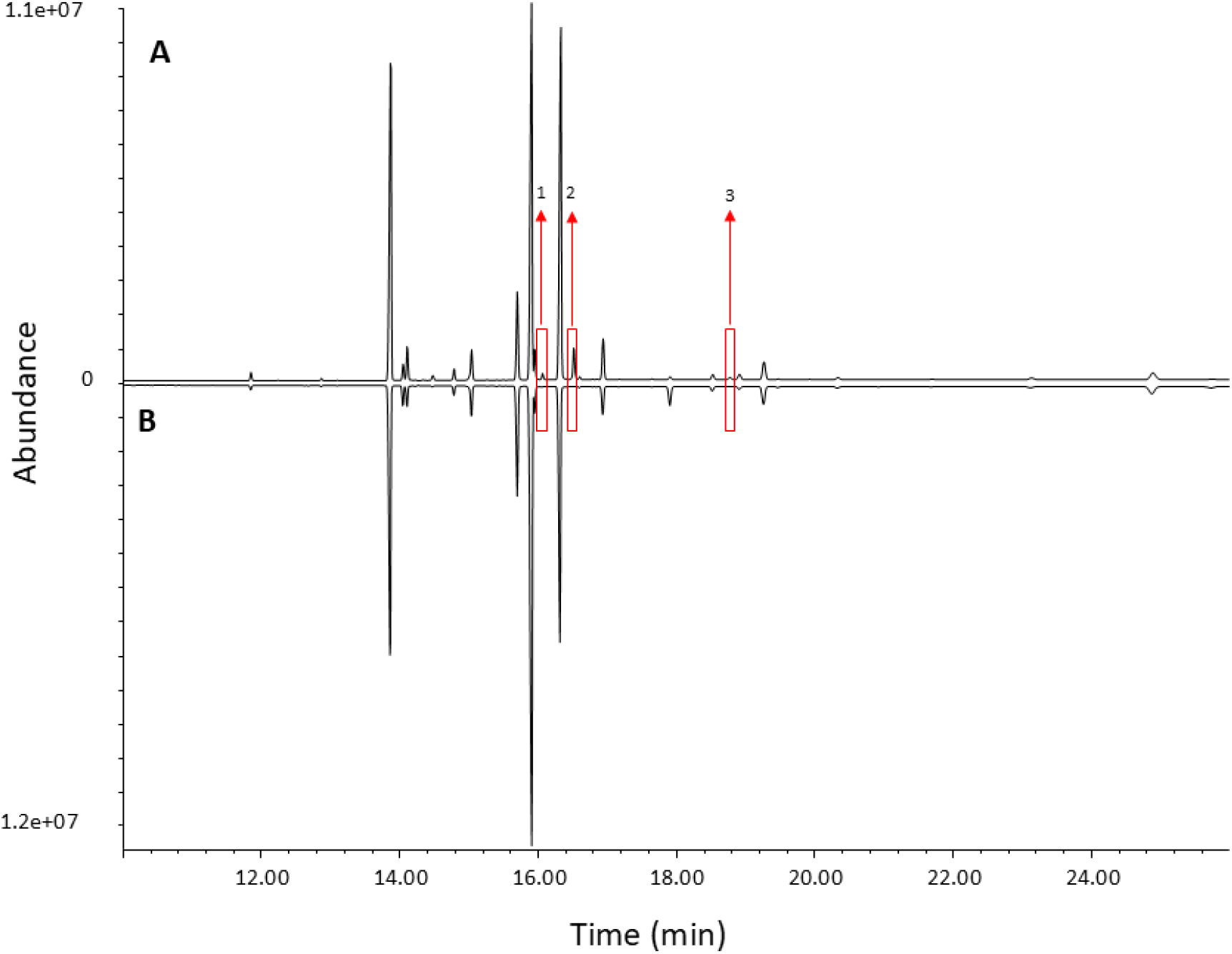
Example of a profile comparing the abundance of fatty acid yolk components between an egg collected in a year with (A) low and (B) high availability of beech (Fagus sylvatica) seeds (year 2016 and 2015, respectively). The red box indicates three fatty acids (1: taxoleic acid - 18:2n-9; 2: pinolenic acid - 18:3n6; and 3: sciadonic acid - 20:3n6) that were present only in the year when beech fructification was low.

## Acknowledgements

We are grateful to Nico Adreani, Sabine Jörg, and Natalia Pérez-Ruiz for their invaluable help in the field. We also thank Martin Laußer for providing us information on the tree composition of the Dellinger Buchet forest, and Dr. Darius Kavaliauskas from the Bavarian Office for Forest Seeding and Planting (ASP) for giving us the seeds used in this study and information of Beech fructification in Bavaria (Germany). We are also grateful to Prof. Dr. Mark van Kleunen and Dr. Veit Martin Dörken for their help and botanical advice.

## Authors’ contributions

Conceptualization: LM and MNA. Methodology: all authors. Software: LM. Validation: LM, HLW and MNA. Formal analysis: LM. Investigation: LM, HLW, MNA, and CI. Resources: MH and CI. Data curation: LM and MNA. Visualisation: LM. Writing - original draft: LM, MNA and CI. Writing - review and editing: all authors. Supervision: LM and MH. Project administration: LM, MH, MNA, and CI. Funding acquisition: MH.

## Ethics statement

All field procedures were conducted in compliance with the legal requirements of Germany and the EU and were approved by the regional governmental authority, the Regierungspräsidium von Oberbayern, Germany (license no. 55.2-1-54-2532-25-2015).

## Data archiving statement

The dataset and the R code generated for this study are available in the supplementary material. Upon acceptance of the paper, they will be openly available in GitHub.

